# Analysis of astrocyte progenitors derived from human induced pluripotent stem cells *in vitro* and following transplantation into the intact spinal cord

**DOI:** 10.1101/2025.06.27.661961

**Authors:** Alessia Niceforo, Ying Jin, Skandha Ramakrishnan, Maurice Lindner-Jackson, Liang O. Qiang, Itzhak Fischer

## Abstract

Functional improvement following traumatic spinal cord injury (SCI) remains limited, therefore, it is necessary to develop therapeutic interventions such as cell transplantation to replace lost cells and promote connectivity. While transplantation typically focuses on neurons, it is important to include other neural cells, such as immature astrocytes, to provide a permissive environment, promote neuroprotection and regeneration, and ultimately restore connectivity. In this study, we leveraged cellular engineering using human induced pluripotent stem cells (hiPSCs) to generate astrocyte progenitor cells (hAPCs). We tested two hiPSC lines (WTC11 and KOLF2.1J) to characterize the fate of the hAPCs *in vitro* and following transplantation at the cervical level of the intact spinal cord for up to 3 weeks. Our results demonstrated efficient and consistent differentiation of the hiPSCs into hAPCs, their survival and integration with the adult spinal cord, with no signs of tumors, deleterious outcomes, and unexpected locations. The ability to survive and the absence of adverse effects indicate that hAPC transplantation could be a safe element of therapy in treating spinal cord injuries.

## INTRODUCTION

Traumatic spinal cord injuries (SCIs) at the cervical level result in an array of deficits, including life-threatening impairments in breathing, due to the disruption of phrenic motor networks located in the cervical spinal cord ^1–4^. The extent of long-term functional improvement associated with spontaneous plasticity following SCI remains limited due to the limited capacity of the adult spinal cord to regenerate, the inhibitory environment of the injury site, and persistent secondary injury^5,6^.

Considering the extensive tissue damage from SCIs, therapeutic treatments need to address cell replacement related to anatomical repair as well as the promotion of regeneration to restore connectivity. In recent years, progress in understanding endogenous plasticity and mechanisms of regeneration has expanded the range of tools and approaches designed to enhance recovery^3,7^.

Cell transplantation has become a promising strategy because it offers multiple molecular and cellular benefits ^6,8,9^. Various transplantation approaches have been explored for SCI, including neuronal and non-neuronal cell types ^10,11^. In the last 2 decades, our labs ^9,11–15^ and others^16,17^ have shown that the transplantation of neural progenitor cells (NPCs) provides the essential neural components for generating new neurons and glial cells (i.e. oligodendrocytes and astrocytes), which are crucial for building neural networks and enhancing connectivity. Neurons derived from NPC transplantation have been shown to integrate both anatomically and functionally with host neural circuits, facilitating the formation of new neuronal relays across the injury site^14,17–19^. Additionally, NPC-derived glial cells offer additional therapeutic advantages in creating a supportive environment for the survival and differentiation of neurons, promoting axon regeneration and remyelination in the formation of functional relays ^11,20–23^ and reducing glial scar formation^21,24^. Although NPC transplantation has demonstrated therapeutic benefits in both preclinical and clinical studies, the extent of recovery following SCI remains limited and inconsistent ^3,5^. It is evident that donor tissues contain a highly heterogeneous population of neural cell types, including neuronal and glial, which may contribute to the variability in outcomes ^25^ and the need for immunosuppression when using allografts present challenges for effective clinical translation. There is a need to test and optimize readily available human cells, preferably with the potential for autologous grafting.

While a complete understanding of the optimal cell types for spinal cord repair is still lacking, recent advancements in stem cell-based therapy offer a promising alternative to address these challenges ^2,26,27^. Understanding the fundamental properties and mechanisms of the different types of stem cells is essential for their therapeutic use in SCI ^28–30^ and other CNS injuries. Among stem cells with the potential to differentiate into all three embryonic germ layers, human induced pluripotent stem cells (hiPSCs)^29,31–33^ provide considerable advantages either as stable cell lines or patient-derived cells. HiPSCs are generated by reprogramming adult somatic cells (i.e. fibroblasts, hematopoietic cells)^34^ to a pluripotent state ^31,35–37^. They have the potential to be ‘patient-specific’, thus avoiding the critical challenge of tissue rejection and/or the use of immunosuppression related to transplant procedures^27,38–40^. Therefore, this approach brings new therapeutic avenues to SCI patients, who currently have few treatment options, making them a crucial resource in regenerative medicine ^27,38,39,41^. In the last decade, researchers have been using the hiPSC-derived neural cells to replace lost neurons, promote axonal regeneration, and provide neurotrophic support, contributing to functional recovery in animal models of SCI ^42–46^. Furthermore, efforts are underway to guide the directed differentiation of iPSCs into specific neural cell types and enhance their integration into host tissue, aiming to maximize their therapeutic potential for SCI treatment ^47,48^.

Here, we show the successful and consistent generation of hAPCs using two hiPSC lines (WTC11 and KOLF2.1J) by relatively simple differentiation protocols. Their survival and phenotype were then tested *in vivo* by transplanting them into the intact cervical spinal cord of rats for up to 3 weeks. We observed that the grafted cells exhibited a stable glial fate at 1- and 3-week post-transplantation, accompanied by survival, morphological maturation, and limited distribution at the injection sites. Importantly, the grafts did not cause any evident adverse effects, and cells remained confined to appropriate locations. The long-term outcomes observed in this study suggest that transplanting specific populations of hAPCs derived from hiPSCs could be a crucial component of the therapeutic approach for cervical SCI treatment.

## MATERIALS AND METHODS

### *In vitro* differentiation of hiPSC cell lines

We purchased two hiPSC lines: WTC11 (Coriell Institute, https://catalog.coriell.org/0/Sections/Search/Sample_Detail.aspx?Ref=GM25256&Product=CC), and KOLF2.1J (Jackson lab, https://www.jax.org/jax-mice-and-services/ipsc).

We differentiated the two hiPSC lines (WTC11 and KOLF2.1J) using the protocol of Stoklund-Dittlau et al. 2023 ^40^. Briefly, the hiPSC lines were maintained on Matrigel-coated plates (Corning; 356231) in mTeSR1™ culture media (Stem Cell Technologies, 85850). The media was changed daily, and hiPSCs were dissociated with ReLeSR™ (Stem Cell Technologies, 100-0484), and the cell clusters were collected in Corning® ultra-low attachment 6-well (Corning, 3471) to support embryoid body (EB) formation. EBs were kept in neuronal induction medium (NIM) consisting of Complete Essential 8™ medium (Thermo Fisher, A1517001) with 1% penicillin/streptomycin (Thermo Fisher, 15140122) supplemented with 0.1 μM LDN-193189 (Reprocell, 04–0074-02) and 10 μM SB431542 (Tocris Bioscience, 1614) for the first week of differentiation. Media changes were performed on days 1, 2, and 4. On day 7, the EBs were plated on Matrigel-coated plates for neural rosette formation and subsequent neural progenitor cell (NPC) expansion in neuronal maturation medium (NMM) containing 50% DMEM/F12 (Thermo Fisher, 11330032) and 50% Neurobasal medium (Thermo Fisher, 21103049) with 1% L-glutamine (Thermo Fisher, 25030081), 1% penicillin/streptomycin, 1% N-2 supplement (Thermo Fisher, 17502048) and 2% B-27™ without vitamin A (Thermo Fisher, 12587–010) supplemented with 10 ng/ml recombinant murine fibroblast growth factor (FGF)-2 (PeproTech, 450–33), 10 ng/ml recombinant human epidermal growth factor (EGF) (ProSpec, Rotherham, CYT-217), 0.1 μM LDN-193189 and 10 μM SB431542. The medium was changed daily, and NPCs were passaged a few times using Accutase (Sigma-Aldrich, A6964). On day 16, NPCs were cultured until day 25 in astrocyte differentiation medium (ADM) made of 90% Neurobasal medium, 1% penicillin/ streptomycin, 1% N-2 supplement, 1% non-essential amino acids (Thermo Fisher, 11140050) and 0.8 μM ascorbic acid (Sigma-Aldrich, A4403) supplemented with 10 ng/ml FGF-2, 200 ng/ml recombinant human insulin-like growth factor (IGF)-1 (Peprotech, 100–11), 10 ng/ml human Activin A (Thermo Fisher, PHC9564) and 10 ng/ ml recombinant human Heregulinβ1 (Peprotech, 100–03) to convert NPCs to astrocyte progenitor cells (hAPCs). Medium changes were made every second day during the period. On day 25, a glial switch is expected to occur, which commences the astrocyte maturation period. APCs were plated for final experiments on new Geltrex®-coated plates and matured for an additional 4 weeks in astrocyte maturation medium (AMM) consisting of 50% DMEM/F12, 50% Neurobasal medium, 1% non-essential amino acids, 1% N-2 supplement, 1% L-glutamine, 1% penicillin/streptomycin, 2% fetal bovine serum (FBS) (R&D Systems, S11150), 0.8 μM ascorbic acid, and 1% sodium pyruvate (Thermo Fisher, 11360–070) supplemented with 200 ng/ml IGF-1, 10 ng/ml Activin A and 10 ng/ml Heregulinβ1. The cells were cryopreserved in 60% FBS, 10% AMM media, and 10% DMSO (Sigma-Aldrich, C6295-50ML) for later use or fixed for the *in vitro* characterization.

### Immunocytochemistry of fixed cells

Culture cells were fixed with 4% paraformaldehyde (PFA) for 15 min, then washed 3 times with phosphate-buffered saline (PBS) 1X for 5 minutes. Cells were then quenched using the quenching solution, consisting of 30% methanol (Sigma-Aldrich, 646377-4L) and 2% hydrogen peroxide solution (H_2_O_2_) (EMD Millipore, HX06353) diluted in PBS 1X for 1 hour at room temperature and then blocked with 10% Normal Donkey Serum (GeneTex, GTX73205), 0.3%Triton X-100 (Fischer Scientific, TX1568-1) in PBS 1X for 2 hours at room temperature. Cells were then incubated with primary antibodies (Table 1) overnight at 4°C. The following day, cells were washed in PBS 1X (3 times for 5 minutes) and incubated with secondary antibodies (Table 2) for 2 hours at room temperature. Immunolabelled cells were then washed 3 times for 5 minutes with PBS 1X, and stained with 4,6-diamino-2-phenylindole (DAPI) (1:10000; Invitrogen, D1306) in PBS 1X for 10 minutes. Glass coverslips (Thermo Scientific, 3323) were mounted on top of the slides using Fluoro-Gel with Tris Buffer (Electron Microscopy Sciences, 17985–10). Cells were imaged and examined using a Leica TCS SP8 with DMi8 inverted optical stand.

**Table 1.**
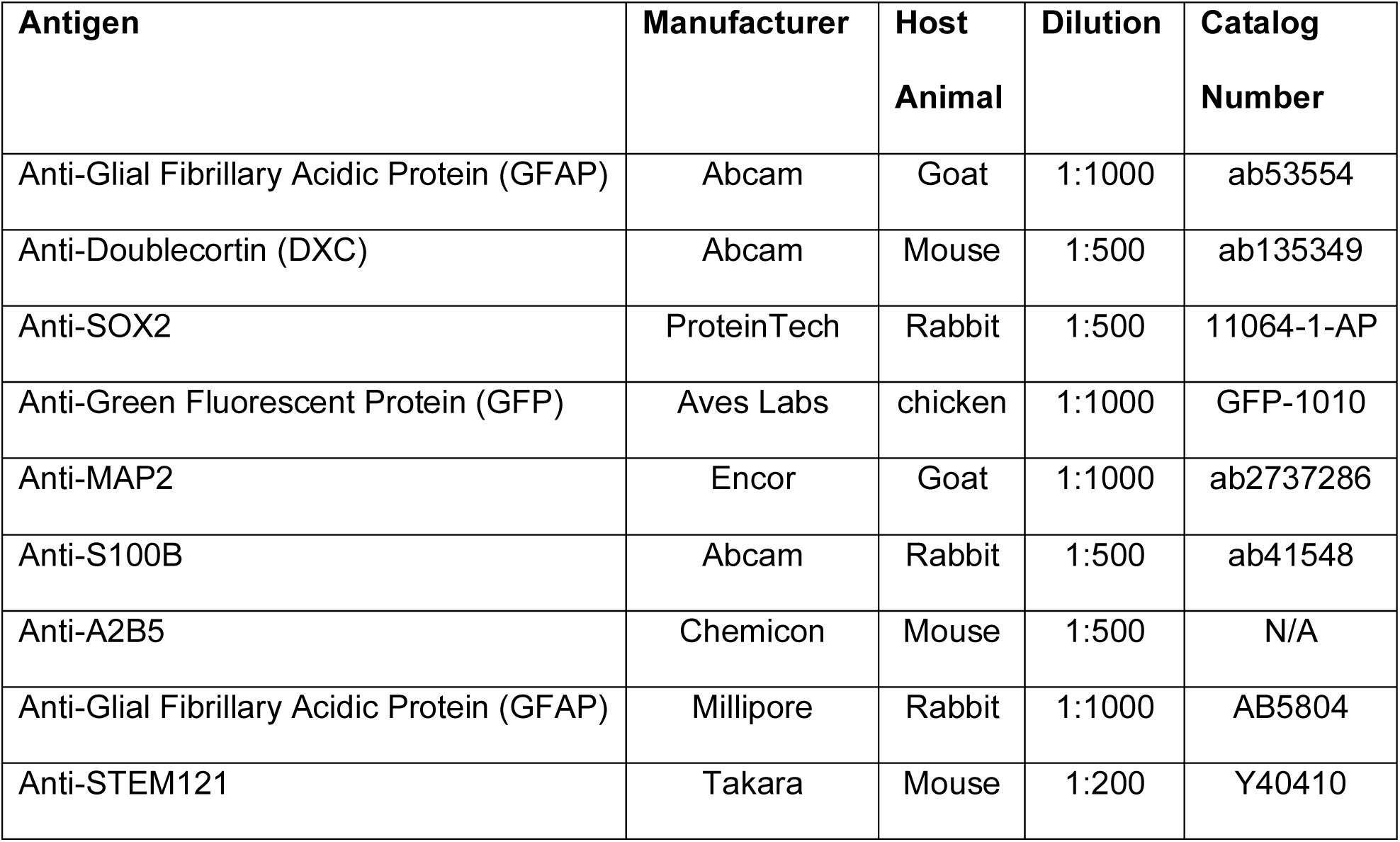
Primary antibodies.

| **Antigen** | **Manufacturer** | **Host**<br>**Animal** | **Dilution** | **Catalog**<br>**Number** |
| --- | --- | --- | --- | --- |
| Anti-Glial Fibrillary Acidic Protein (GFAP) | Abcam | Goat | 1:1000 | ab53554 |
| Anti-Doublecortin (DXC) | Abcam | Mouse | 1:500 | ab135349 |
| Anti-SOX2 | ProteinTech | Rabbit | 1:500 | 11064-1-AP |
| Anti-Green Fluorescent Protein (GFP) | Aves Labs | chicken | 1:1000 | GFP-1010 |
| Anti-MAP2 | Encor | Goat | 1:1000 | ab2737286 |
| Anti-S100B | Abcam | Rabbit | 1:500 | ab41548 |
| Anti-A2B5 | Chemicon | Mouse | 1:500 | N/A |
| Anti-Glial Fibrillary Acidic Protein (GFAP) | Millipore | Rabbit | 1:1000 | AB5804 |
| Anti-STEM121 | Takara | Mouse | 1:200 | Y40410 |

**Table 2.** Secondary antibodies.

| **Antigen**<br>**Animal** | **Manufacturer** | **Host Animal** | **Fluorophore** | **Dilution** | **Catalog**<br>**Number** |
| --- | --- | --- | --- | --- | --- |
| Chicken-IgG | Jackson Immuno<br>Research | Goat | 488 | 1:1000 | 103-295-155 |
| Goat-IgG | Jackson Immuno<br>Research | Donkey | 647 | 1:500 | 705-605-003 |
| Mouse-IgG | Invitrogen | Donkey | 488 | 1:500 | A21202 |
| Mouse-IgM | Jackson Immuno<br>Research | Donkey | 647 | 1:500 | 715-605-020 |
| Mouse-IgG | Invitrogen | Donkey | 647 | 1:500 | A31571 |
| Rabbit-IgG | Jackson Immuno<br>Research | Donkey | 594 | 1:500 | 711-585-152 |

All images from each replicate were acquired with identical exposure settings. The fluorescence intensity of each channel was quantified using the ImageJ software and normalized per the number of nuclei (DAPI-positive cells).

### Quantitative Real-Time Polymerase Chain Reaction (qRT-PCR)

The RNA extracted from cultured cells was isolated using the “PureLink™ RNA Mini Kit” (Invitrogen™, 12183018A) and reverse transcribed into cDNA using the “High-Capacity cDNA Reverse Transcription Kit’ (Applied Biosystems™, 4368814). The PCR reaction was carried out with the “2X Universal SYBR Green Fast qPCR Mix” (ABclonal, RM21203) using the “7900 Real-Time PCR system” (Applied Biosystems). The primers used are listed in Table 3. Gene expression was analyzed using the 2-Delta-Delta Ct method. All results were normalized to GAPDH expression. Three technical replicates per run and three biological replicates for each sample were used to determine the error bars.

**Table 3.**
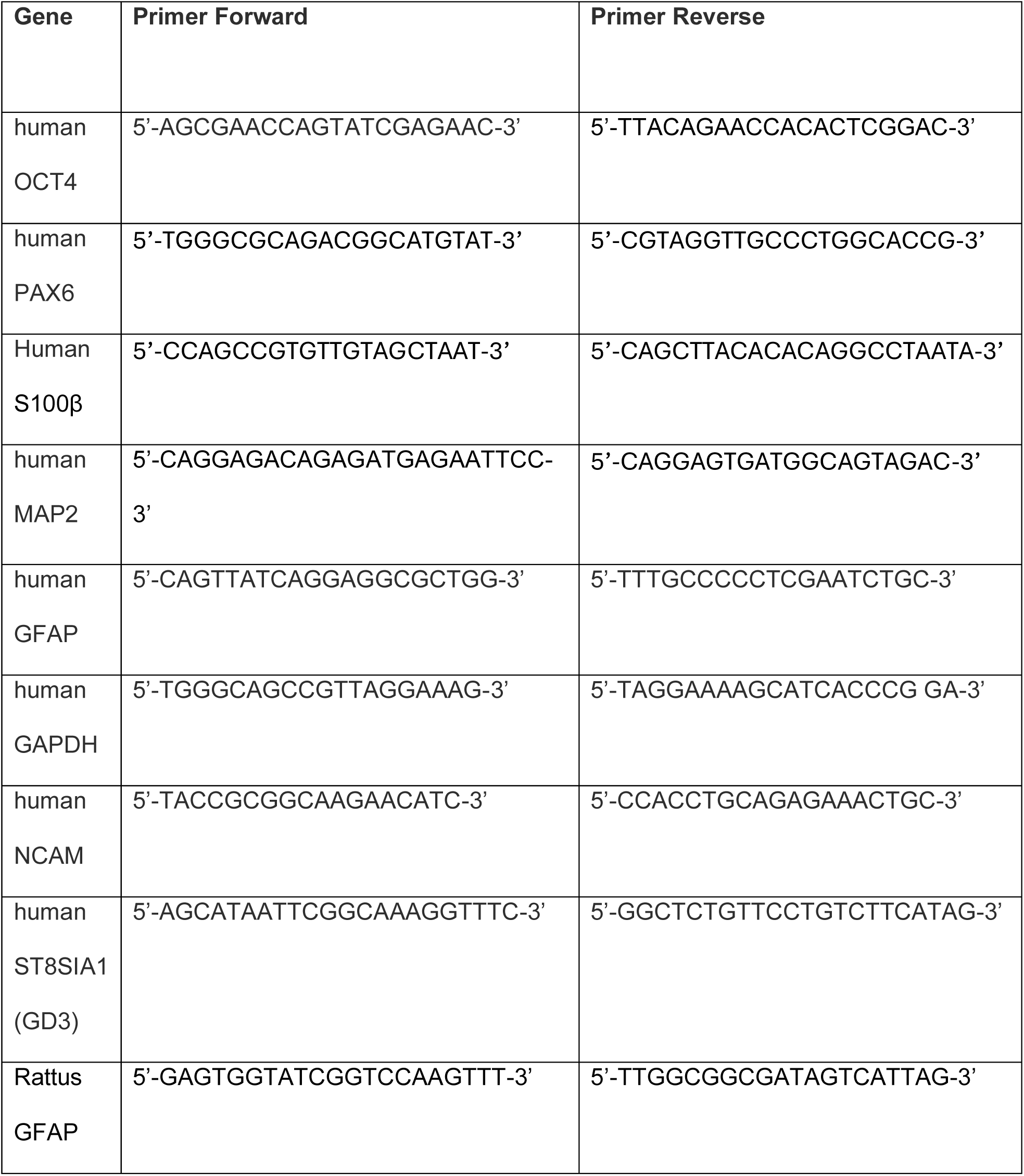
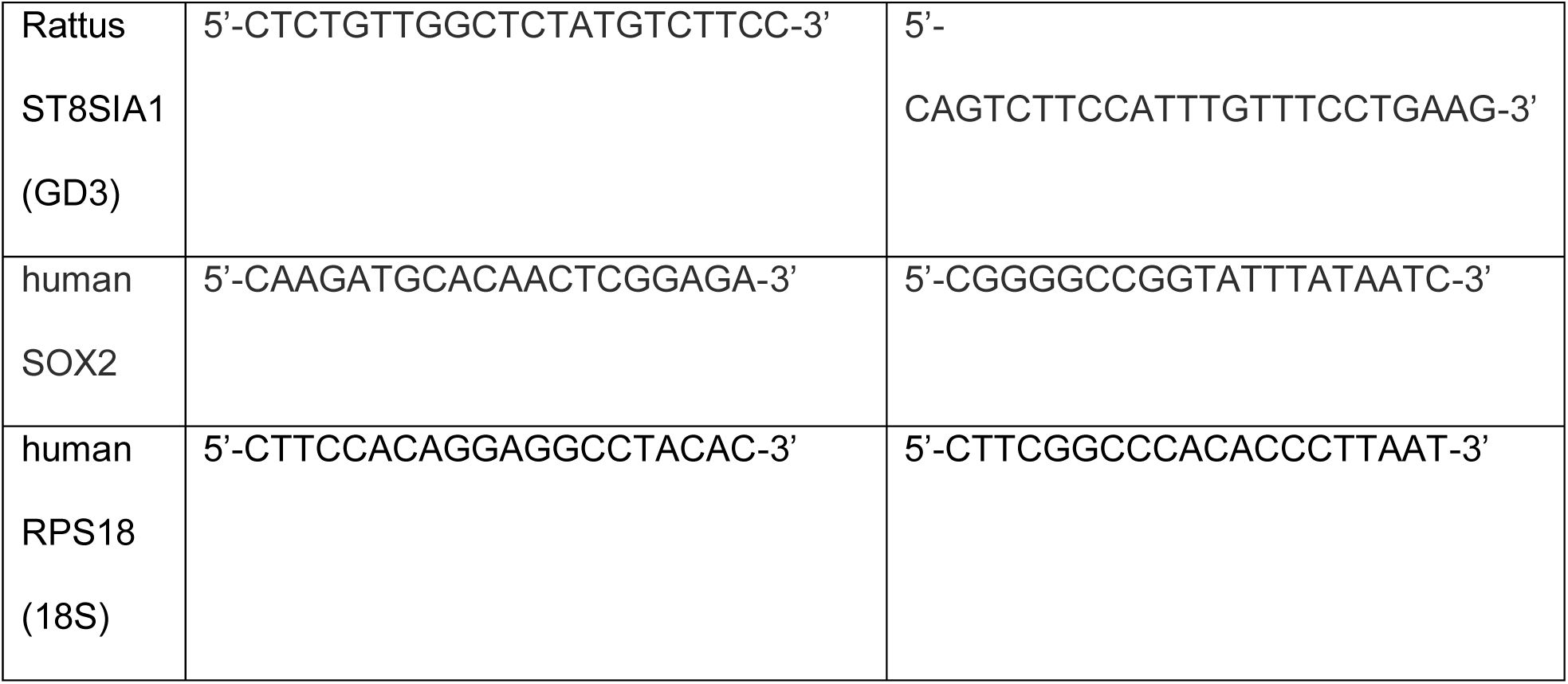
Primers for quantitative real-time (qRT-PCR)

### Statistical Analysis

All data analyses were conducted blind and presented as mean ± SEM. Three biologically independent repeats were conducted for each experiment. Data were arranged in Microsoft Excel before statistical analyses were performed using Prism 10 (GraphPad). The statistical tests used were one-way analysis of variance (ANOVA). The Kruskal–Wallis test, followed by Dunn’s multiple comparison test, was used for data not normally distributed. Statistical significance was defined as *p<0.05.

### Green Fluorescent Protein (GFP^+^) lentiviral production

For our experiments, we bought the plasmid *pLenti CMV GFP Hygro (656-4)* from AddGene (plasmid #17446) (https://www.addgene.org/17446/). The lentivirus GFP^+^ was produced as follows: a vial of 293FT cells (obtained from our collaborator Dr. Qiang), was thawed onto 100 mm TC-treated culture dishes (Corning, 430167) and maintained in Dulbecco’s Modified Eagle’s Medium (DMEM) (Fisher Scientific, 10-566-016) supplemented with 10% FBS. Once the cells reached 80% confluency, the GFP^+^ plasmid was packaged into a lentiviral vector using second-generation packaging plasmids and the calcium chloride precipitation method of transfection. Briefly, on the day of transfection, 293FT cells were co-transfected with GFP, pMD2.G (Addgene, 12259), and psPAX2 (Addgene, 12260) plasmids. After 48 hours, the virus-containing medium was collected, centrifuged, and filtered using a 0.45μm filter. The virus was then concentrated using Lenti-X™ Concentrator (Takara Bio, 631232), aliquoted, and stored at −80 °C according to the manufacturer’s instructions.

### hAPC infection and preparation for engraftment

48 hours before the grafting, the WTC11- hAPCs and KOLF2.1J- hAPCs were infected using 2.5µl of lentivirus *GFP^+^* (10^7^ - 10^8^ pfu) in AMM media. The day later, half of the media was changed with fresh media. 1 hour before the grafting, WTC11-hAPCs-GFP^+^ and KOLF2.1J- hAPCs-GFP^+^ were washed in PBS 1X, and lifted using accutase for 5 min at 37°C. Cells were collected and counted using the automated cell counter “Cellometer Auto T4 (Nexcelom Bioscience). The cells were then resuspended in Hank’s Balanced Salt Solution1X (HBSS) (Gibco, 24020-117) before grafting.

### Animal Surgery/Graphing

Adult, female, Sprague Dawley rats (n=6) (approximately 200g) were housed at the Drexel University College of Medicine animal care facility. All experimental procedures were conducted with approval from the Institutional Animal Care and Use Committee and following the National Research Council Guidelines for the Care and Use of Laboratory Animals. Animals received the transplanted cells into the grey matter of the intact spinal cord during the same grafting session. Three days before the cell grafting, all the animals received subcutaneous (s. q.) injections of the immunosuppressive drug Cyclosporine A (CSA) (10mg/kg) (McKesson, 1622448). The CSA was delivered to the animals daily until the end of the experiments (7- or 21-day post-grafting).

On the day of the surgery, animals were anesthetized using isoflurane (4% in O_2_ to induce, 2% in O_2_ to maintain). The back of the neck (from the base of the skull to the top of the shoulder blades) was shaved and sterilized with alternating alcohol 70% and iodine solution (ATC Medical, 233). An approximately 2.5-inch skin incision was made with a No. 15 scalpel blade from the base of the skull to the fifth cervical segment (C5), and the surrounding musculature was cut and carefully pushed apart to expose the vertebral column. A laminectomy was made at the third cervical segment (C3) and the rostral part of C4. Animals were blindly separated into one of two groups (WTC11-hAPCs-GFP^+^, KOLF2.1J-hAPCs-GFP^+^) and decoded after all data analysis.

The surgery was conducted as previously done in our lab ^16^. Briefly, each animal received two grafts of 10,000 cells/each, resuspended in 2 μl of HBSS at the C3-4 level.

Cells were injected into the grey matter of the intact spinal cord using a 10μL Hamilton syringe (Hamilton, 84853) with a custom 30-gauge needle attached to a micromanipulator. One animal was injected with 10,000 of WTC11-hAPCs resuspended in 2 μl of HBSS at the C3-4 level as a negative control. One animal died immediately after the grafting.

Following surgeries, the underlying muscle was sutured using 4-0 Vicryl Sutures (Fischer Scientific, VCP310H) in layers, and the skin was closed with wound clips (9mm; Fischer Scientific, NC0142166). Animals received lactated ringers (5 mL, s.q.) to prevent dehydration and buprenorphine (0.025mg/kg, s.q.) as an analgesic.

### Tissue processing

Animals were sacrificed at 2 time points (7- and 21-day post-grafting) by intracardial perfusion with physiological saline (0.9% NaCl in water), followed by ice-cold 4% PFA (in 0.1 M PBS; pH 7.4). The spinal cords were removed from the animals and stored in 4% PFA at 4°C overnight, followed by cryoprotection in 30% sucrose/0.1 M PBS at 4°C for 3 days. The tissues were then embedded in M1 Embedding Matrix (Thermo Fischer, 1310TS), fast-frozen with dry ice, and sectioned on-slide (40μm thickness, transverse) using a cryostat. Sections were stored at −20 °C until analyzed.

### Immunocytochemistry of tissues

To perform immunohistochemistry (IHC), sections were rehydrated by washing in PBS 1X (3 x 10 min) and incubated for 2 hours with the blocking solution consisting of 25 mM Tris HCl, 300 mM NaCl, 0.3%Triton X-100, 0,5 mg/mL bovine serum albumin (BSA, Sigma Aldrich, A4737), 0.01% Thimerosal (Sigma, T-5125), and 10% Normal Goat Serum. The solution was then removed, and the primary antibodies (Table 1) were incubated overnight at 4°C. The following day, tissues were washed in PBS 1X (3 × 10 min) and incubated with secondary antibodies (Table 2). Immunolabeled sections were then washed in PBS 1X (3 × 10 min), allowed to dry for 30 min, and mounted with DAPI using glass coverslips. Sections were examined using a Leica TCS SP8 with DMi8 inverted optical stand.

Cell survival at 7- and 21-day post-transplantation was quantified by counting the total number of hAPCs-GFP^+^ cells positive for the staining hSTEM121 in the grey matter of the spinal cord, and dividing by the initial number of cells injected (10,000 cells).

## RESULTS

### *In vitro* differentiation and characterization of hAPCs

We used two induced pluripotent stem cell (iPSC) lines, WTC11 and KOLF2.1J, to validate the consistency of the differentiation protocols. These hiPSC lines were selected based on recent studies demonstrating their strong potential for differentiation into glial progenitor cells ^40,49–52^. We differentiated the two hiPSC lines (WTC11 and KOLF2.1J) using the protocol of Stoklund-Dittlau et al. 2023 ^40^.

The *in vitro* differentiation of hiPSCs into astrocytes became evident after 35+ days, using specific growth factors and media ^40^ as shown in Fig.1A. At the start of the differentiation process (day 0), phase-contrast images revealed that the WTC11 iPSCs and the KOLF2.1J hiPSCs exhibited the typical morphology of colonies, characterized by round shape, defined border, and uniformity (Fig. 1B a, a’). As differentiation progressed, the cells underwent distinct morphological transitions as follows:

**Figure 1:**
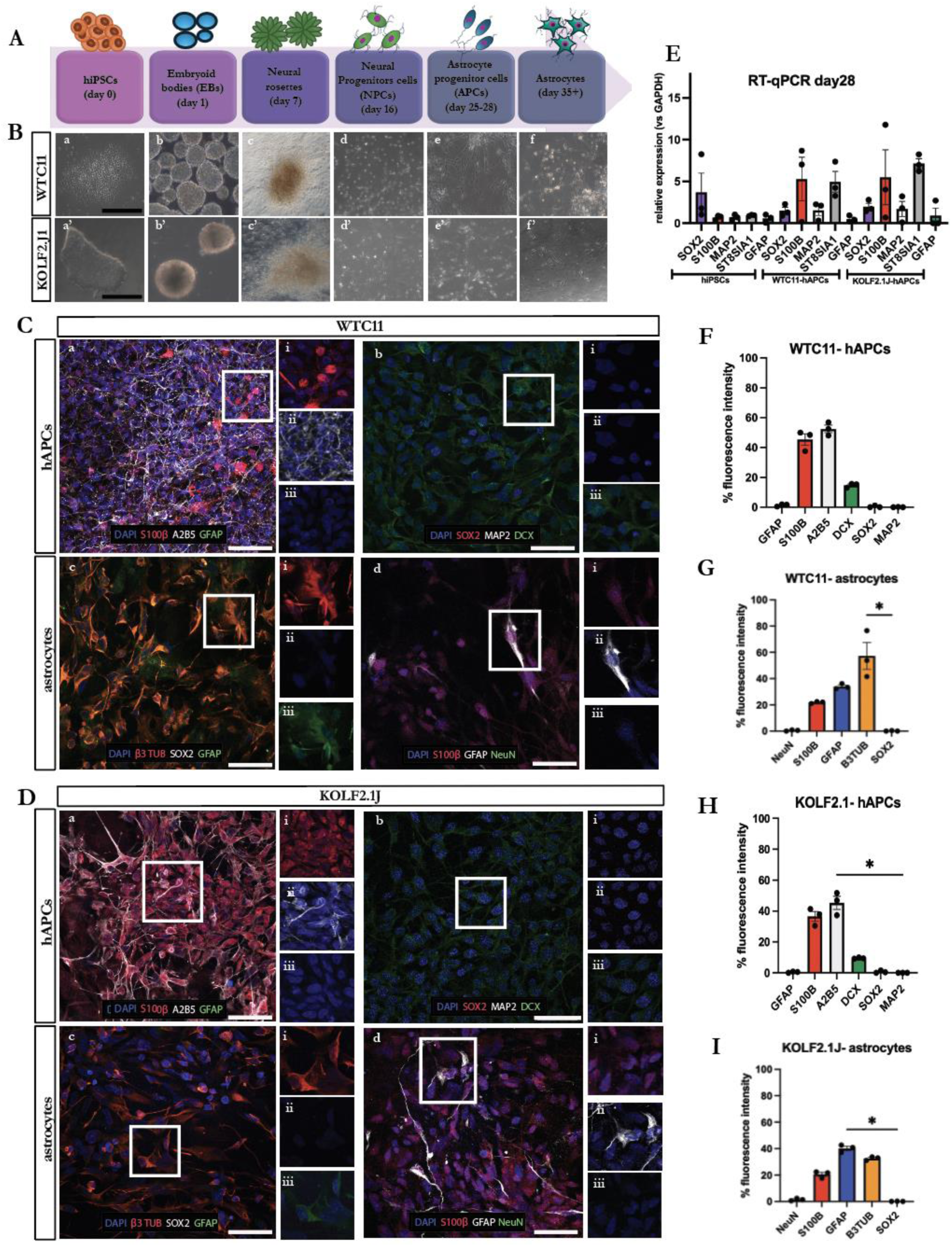
(A) A schematic of the experimental timeline showing the differentiation steps of hiPSCs (day 0) to astrocytes (day 35), going through the phases: EBs (day 1), neural rosette (day 7), NPCs (day 16), APCs (day 24-28). (B) During astrocyte differentiation, phase-contrast pictures reveal changes in the morphology of the 2 hiPSC lines WTC11 (a-f), and KOLF2.1J (a’-f’). Scale bar: 250 microns in a-f and a’-f’. (C) Immunocytochemistry (ICC) of WTC11-derived APCs (day 28), and WTC11-derived astrocytes (day 35). The markers used were the astrocytic marker glial fibrillary acidic protein (GFAP), neuronal nuclear protein NeuN, the astrocyte progenitor marker S100β, neuronal precursor marker doublecortin (DCX), the cytoskeletal marker β3-tubulin (B3TUB), neural stem cells marker SOX2, and glial-restricted progenitor marker A2B5. Nuclei were stained blue with DAPI. Scale bar: 20 microns in C a-d. C ai-iii: magnification of Ca; C bi-iii: magnification of Cb; C ci-iii: magnification of Cc; C di-iii: magnification of Cd. (D) ICC of the KOLF2.J1-derived cells at 2 different stages: astrocyte progenitor cells APCs (day 28) and astrocytes (day 35). Scale bar: 20 microns in D a-d. D ai-iii: magnification of Da; D bi-iii: magnification of Db; D ci-iii: magnification of Dc; D di-iii: magnification of Dd. (E) RT-qPCR showing the different gene expressions of hiPSCs, WTC11-derived APCs, and KOLF2.1J-derived APCs at day 28 of differentiation. The *GAPDH* was used as the housekeeping gene. Data are presented as mean ± SEM of 3 independent experiments. Analyses were conducted using one-way ANOVA and the Kruskal-Wallis test. (F-I) Analyses of fluorescence intensity at day 28 of differentiation (F-H) confirmed that WTC11-derived APCs, and KOLF2.1J-derived APCs were positive for the glial-restricted progenitor marker A2B5, doublecortin (DCX), and S100β. (G-I) Astrocytes at day 32 tested positive for the astrocytic marker GFAP, β3-tubulin (B3TUB), and S100β. Data are represented as mean ± SEM of 3 independent experiments. Analyses were conducted using one-way ANOVA and the Kruskal-Wallis test. *p<0.05.

Day 1: Cells formed aggregates in suspension as embryoid bodies (EBs) (Fig. 1B b, b’).

Day 7: Cells radially arranged around a central lumen, forming neural rosettes (Fig. 1B c, c’).

Day 16: At the neural progenitor cell (NPC) stage, cells transitioned from aggregates to a spread, single-cell organization (Fig. 1B d, d’).

Days 25–28: During the astrocyte progenitor cell (APC) stage, cells exhibited small nuclei and elongated cell bodies (Fig. 1B e, e’).

Day 35: Cells began adopting a star-shaped phenotype, characteristic of mature astrocytes (Fig. 1B f, f’). These morphological changes confirm the successful differentiation of hiPSCs into astrocyte progenitors and eventually into mature astrocytes.

To assess the detailed composition of the hiPSC-derived astrocyte progenitor cells (hAPCs, day 28) and hiPSC-derived astrocytes (day 35), analyses of immunocytochemistry (ICC) were conducted (Fig. 1C, 1D). Our data showed that the hAPCs (day 28) obtained from both the two hiPSC lines (WTC11 and KOLF2.1J) were positive for the glial-restricted progenitor marker A2B5, the astrocyte progenitor marker S100β, and the neuronal progenitor marker doublecortin (DCX), indicating the glial progenitor status of the majority of these cells. Additionally, the WTC11-hAPCs and the KOLF2.1J-hAPCs tested negative for the neural stem cell marker SOX2, and the neuronal marker MAP2, confirming the absence of undifferentiated stem cells and neurons, respectively (Fig. 1C a-b, 1D a-b). At day 35, hiPSC-derived astrocytes from both cell lines, tested positive for Glial fibrillary acidic protein (GFAP), showing a star-shaped morphology typical of mature astrocytes. However, some cells tested positive for β3-tubulin (β3 TUB), and the astrocyte progenitor marker S100β (Fig. 1C c-d, 1D c-d), indicating their persistence in the state of precursor astrocytes.

Quantitative analysis of fluorescence intensity reinforced these findings (Fig. 1F-I). At day 28 of differentiation, the WTC11 -hAPCs and KOLF2.1J -hAPCs were positive for the markers A2B5 (WTC11- hAPCs 52%, KOLF2.1J -hAPCs 45%), DCX (WTC11- hAPCs 14%, KOLF2.1J -hAPCs 9%), and S100β (WTC11- hAPCs 45%, KOLF2.1J -hAPCs 36%) (Fig 1 F-H). Notably, the GFAP marker expression was absent in KOLF2.1J- hAPCs, although it was expressed in a limited number of cells in WTC11- hAPCs at day 28 (2%).

At day 35, the WTC11-astrocytes, and KOLF2.1J- astrocytes tested positive for the markers GFAP (33% and 40%, respectively), the β3-tubulin (WTC11- astrocytes 57%, and KOLF2.1J- astrocytes 32%), and S100β (WTC11- astrocytes 21%, and KOLF2.1J- astrocytes 20%) (Fig 1 G-I).

To further validate the phenotype of the differentiated cells, we performed the gene expression analysis by RT-qPCR of WTC11-hAPCs and KOLF2.1J-hAPCs at day 28, compared them with undifferentiated hiPSCs (Fig. 1E). Our results showed that the hiPSCs expressed *SOX2*, confirming the pluripotency state, although low levels of MAP2, GFAP, and S100β were also detected. RT-qPCR further revealed high expression of the glial-restricted progenitor gene *ST8SIA1,* and of the astrocyte progenitor marker *S100β* gene, in both WTC11-hAPCs and KOLF2.1J-hAPCs. Low expression of the GFAP, MAP2, and SOX2 genes was also detected in KOLF2.1J-derived hAPCs.

These findings confirm the glial lineage commitment of the differentiated cells while highlighting no significant differences in marker expression between the two iPSC lines.

### Grafted WTC11-hAPCs and KOLF2.1J-hAPCs survived 1 week and 3 weeks after transplantation

Forty-eight hours before the grafting (DV −2, Fig. 2A), the WTC11-hAPCs and KOLF2.1J- hAPCs were infected using 2.5µl of lentivirus *GFP^+^* in AMM media. Half of the media was changed with fresh media the day later (DV −1, Fig. 2A). One hour before the grafting, WTC11-hAPCs-GFP^+^ and KOLF2.1J-hAPCs-GFP^+^ were washed in PBS 1X, counted, and then resuspended in HBSS 1X (DV 0, Fig. 2A). The phase-contrast pictures of WTC11-hAPCs-GFP^+^ (Fig. 2B), and KOLF2.1J-hAPCs-GFP^+^ (Fig. 2C) showed that cells were healthy, exhibiting small nuclei and elongated cell bodies on the grafting day (DV 0).

**Figure 2:**
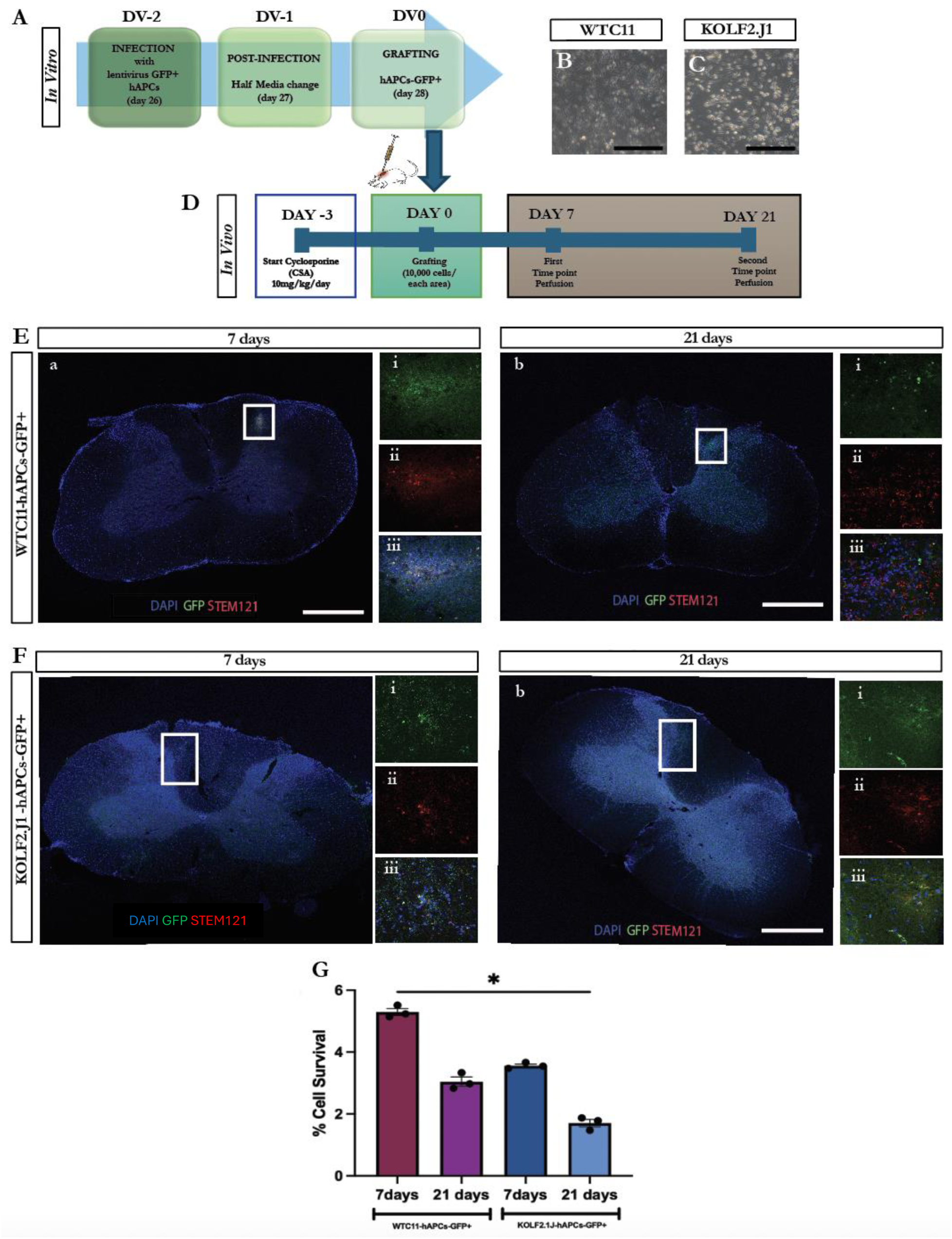
(A) A scheme of the *in vitro* experiment showing the preparation of the hAPCs to be grafted. Both the two hAPCs lines (WTC11-hAPCs and KOLF2.1J- hAPCs) were infected with the lentivirus *GFP^+^* 48 hours before the grafting (DV −2). The day later (DV −1), half of the media was changed with fresh media. (B-C) Phase-contrast pictures at the grafting day (DV0) of hAPCs- GFP^+^ (WTC11-hAPCs and KOLF2.1J-hAPCs, respectively). Scale bar: 250 microns in B and C. (D) A schematic timeline of the *in vivo* experiments. (E) Immunohistochemistry (IHC) of the grafted WTC11-hAPCs-GFP^+^ in the intact spinal cord at 7- (Ea) and 21-day post-grafting. (Eb). E ai-iii: 63X magnification of Ea; E bi-iii: 63X magnification of Eb. Scale bar: 20 microns in E a,b (F) IHC of the grafted KOLF2.1J-hAPCs-GFP^+^ in the intact spinal cord at 7 days (F a), and 21 days (F b) post-grafting. F ai-iii: 63X magnification of Fa; F bi-iii: 63X magnification of Fb. Scale bar: 20 microns in F a,b. The markers used in E and F were Green Fluorescent Protein (GFP) (green) and the human neural stem cell marker hSTEM121 (red). Nuclei were stained blue with DAPI. (G) Cell survival at 7- and 21-day post-transplantation was quantified by counting the total number of hSTEM121^+^ cells in the grey matter of the spinal cord, and dividing by the initial number of cells injected (10,000 cells). Data are represented as mean ± SEM of 3 experiments. Analyses were conducted using one-way ANOVA and the Kruskal-Wallis test. *p<0.05.

For the *in vivo* experiment, 6 adult female, Sprague Dawley rats were used. Three days before the cell grafting (day −3), all the animals received daily s.q. injections of the immunosuppressive drug CSA (10mg/kg), until the end of the experiments (Fig. 2D).

On the day of the surgery (day 0), each animal received grafts of 10,000 cells into the grey matter of the intact spinal cord at the C3-4 level (Fig. 2D and Table 4).

**Table 4.** *In vivo* experimental design.

| **Animals** | **Species** | **Cells**<br>**transplanted** | **Region of**<br>**transplant** | **Immune**<br>**suppression** | **Sacrificing**<br>**Time** |
| --- | --- | --- | --- | --- | --- |
| #1 | Sprague<br>Dawley,<br>adult, female | WTC11-hAPCs<br>(GFP<sup>+</sup>) | Spinal cord<br>(Grey matter) | Yes | Died after cell<br>transplantation |
| #2 | Sprague<br>Dawley,<br>adult, female | WTC11-hAPCs<br>(GFP<sup>+</sup>) | Spinal cord<br>(Grey matter) | Yes | 7 days post-<br>grafting |
| #3 | Sprague<br>Dawley,<br>adult, female | WTC11-hAPCs<br>(GFP<sup>+</sup>) | Spinal cord<br>(Grey matter) | Yes | 21 days post-<br>grafting |
| #4 | Sprague<br>Dawley,<br>adult, female | KOLF2.1J-<br>hAPCs<br>(GFP<sup>+</sup>) | Spinal cord<br>(Grey matter) | Yes | 7 days post-<br>grafting |
| #5 | Sprague<br>Dawley,<br>adult, female | KOLF2.1J-<br>hAPCs<br>(GFP<sup>+</sup>) | Spinal cord<br>(Grey matter) | Yes | 21 days post-<br>grafting |
| #6 | Sprague<br>Dawley,<br>adult, female | WTC11-hAPCs<br>(GFP<sup>-</sup>) | Spinal cord<br>(Grey matter) | Yes | 7 days post-grafting |

To evaluate the WTC11-hAPCs-GFP^+^ and KOLF2.1J-hAPCs-GFP^+^ ability to survive *in vivo* after the transplantation, animals were sacrificed at 2 time points (7- and 21-day post-grafting), and histological assessments of grafted cells were performed (Fig. 2E, F).

On day 7 post-grafting, the immunohistochemistry (IHC) analysis showed that the WTC11- hAPCs-GFP^+^ grafted in the grey matter of the intact spinal cord survived (Fig. 2E a). This can be observed by using the expression of the green fluorescent protein GFP (green), combined with the expression of the human neural stem cell marker STEM121 (red). Nuclei were stained blue with DAPI. The 63X magnification pictures in E ai,ii,iii confirmed the presence of the grafted cells(Fig. 2E ai,ii,iii).

On day 21 post-grafting, the transplanted cells in the intact spinal cord exhibited positive staining for GFP and STEM121, confirming the presence of WTC11-hAPCs-GFP^+^ (Fig. 2E b). Although at 21 days post-grafting, WTC11-hAPCs-GFP^+^ showed a similar localization to the cells at 7 days post-grafting, the phenotype of some cells appeared to have changed. Indeed, high magnification showed cells positively labeled with STEM121 with more elongated shapes (Fig. 2E b i-iii).

A similar result was observed with the KOLF2.1J-hAPCs-GFP^+^, on day 7 and 21 post-grafting (Fig. 2F). Indeed, on day 7 post-grafting, the KOLF2.1J-hAPCs-GFP^+^ cells were identified as positively labeled for GFP and STEM121 (Fig. 2F a), confirming the survival of the grafted cells. The magnification pictures in 2F confirmed that (Fig. 2F ai-iii).

Like WTC11- hAPCs-GFP^+^, the KOLF2.1J-hAPCs-GFP^+^ also exhibited positive staining for GFP and STEM121 on day 21 post-grafting (Fig. 2F b), with a distribution pattern resembling that observed at day 7 post-grafting. Notably, the magnified images revealed STEM121- and GFP-positive cells with more elongated and enlarged shapes, suggesting a potential phenotypic change (Fig. 2F b i-iii).

Evaluation of the number of transplanted cells that survived *in vivo* is shown in Fig. 2G. Quantification of hSTEM121^+^ cells in the spinal cord at 7-, and 21-days post-transplantation showed that the transplanted cells survived for up to 21 days, although survival was limited in both the cell lines (∼5% WTC11 at 7-day; ∼3% WTC11 at 21-day; ∼4% KOLF2.1J WTC11 at 7-day; ∼ 2% KOLF2.1J WTC11 at 21-day) (Fig. 2G). This could be a consequence of limited cell proliferation *in vivo.* Notably, when comparing the number of surviving cells at 7- and 21-day post-grafting within each line, we observed that both WTC11 and KOLF2.1J lost around 50% of the transplanted cells.

### Characterization of the grafted WTC11-hAPCs 1 week and 3 weeks after transplantation

To investigate possible changes in the phenotype, analyses of immunohistochemistry of grafted WTC11-hAPCs-GFP+ were conducted at day 7 (Fig. 3A) and day 21 (Fig. 3B).

**Figure 3:**
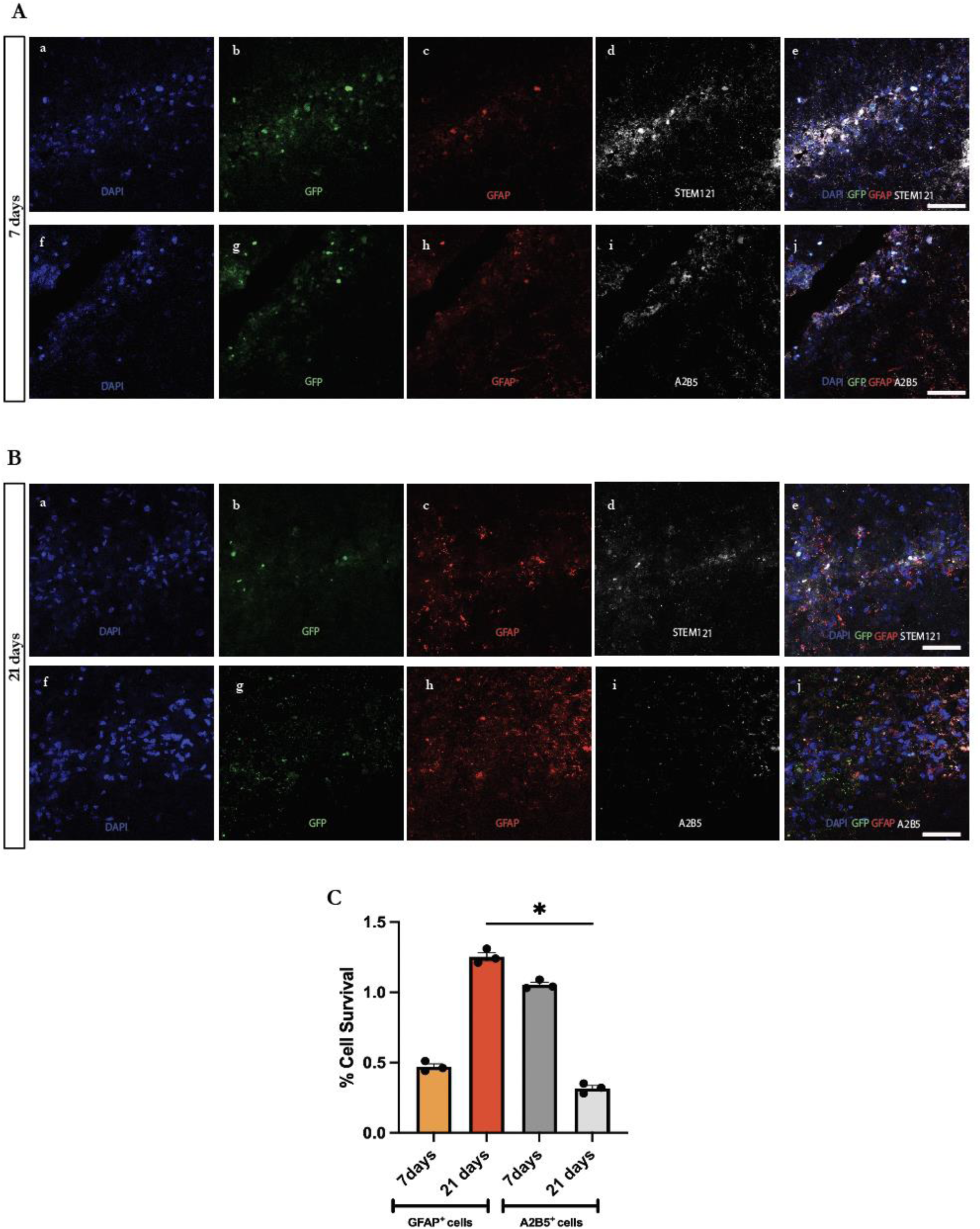
(A) Characterization of the WTC11-hAPCs-GFP^+^ at day 7 post-grafting. The IHC experiments showed that, on day 7, the transplanted cells expressed the green fluorescent protein GFP^+^ in green (b, g), astrocytic marker glial fibrillary acidic protein GFAP in red (c, h), the human neural stem cells marker hSTEM121 in white (d), and glial-restricted progenitor marker in white A2B5 (i). The merged pictures were shown in e and j. Nuclei were stained blue with DAPI (a,f). Scale bar: 20 microns in a-j. (B) Characterization of the WTC11-hAPCs-GFP^+^ at day 21 post-grafting. The markers used were: GFP^+^ in green (b, g), GFAP in red (c, h), hSTEM121 in white (d), and A2B5 in white (i). The merged pictures were shown in e and j. Nuclei were stained in blue with DAPI (a,f). Scale bar: 20 microns in a-j. (C) Cell survival at 7- and 21-day post-transplantation was quantified by counting the total number of double-stained cells hSTEM121-A2B5^+^ and hSTEM121-GFAP^+^ in the grey matter of the spinal cord, and dividing by the initial number of cells injected (10,000 cells). Data are represented as mean ± SEM of 3 experiments. Analyses were conducted using one-way ANOVA and the Kruskal-Wallis test. *p<0.05.

On day 7 post-grafting, the IHC showed that the WTC11-hAPCs grafted in the C3-4 intact spinal cord expressed GFP marker in green (Fig. 3A b, g), combined with the presence of the human neural stem cell marker STEM121 in white (Fig. 3Ad). Noteworthy is the presence of the astrocytic marker GFAP (Fig. 3Ac, red), which indicated a slow and gradual change in the phenotype of the transplanted cells, although the glial-restricted progenitor marker A2B5 was still expressed in some grafted cells (Fig. 3Ai, white). Nuclei were stained blue with DAPI (Fig. 3A a, f).

On day 21 post-grafting, the WTC11-hAPCs-GFP+ also exhibited positive staining for GFP (Fig. 3B b, g), and STEM121 (Fig. 3Bd). Importantly, a notable difference in the expression of GFAP and A2B5 markers was observed. Specifically, the astrocytic marker GFAP showed increased expression at 21 days post-grafting compared to the 7-day time point (Fig. 3Bc, red). In contrast, the progenitor marker A2B5 exhibited low expression at day 21 (Fig. 3Bi, white). These findings, as expected, suggest that the transplanted hAPCs underwent phenotypic changes, differentiating into mature astrocytes. Nuclei were stained blue with DAPI (Fig. 3B a,f).

Quantification of A2B5^+^ cells in the spinal cord at 7- and 21-day post-transplantation revealed that glial-restricted progenitor cells persisted up to 21 days, although their survival was limited (∼%1 at day 7; ∼% 0.3 at day 21) (Fig. 3C). In contrast, the number of mature astrocytes (GFAP^+^ cells) was reduced at 7 days post-transplantation (∼%1.3 at day 7) and increased by 21 days post-transplantation (∼% 0.5 at day 21) (Fig. 3C).

## DISCUSSION

### Therapeutic challenges in spinal cord injury

Traumatic spinal cord injury (SCI), particularly at the cervical level, results in significant neuronal loss, especially affecting the phrenic network, which leads to respiratory dysfunctions ^53–55^. The limited intrinsic regenerative capacity of the central nervous system (CNS) underscores the urgent need for innovative therapeutic strategies to promote functional repair^1,56,57^. Cell-based therapies, specifically those utilizing hiPSC-derived populations, have emerged as promising candidates for addressing these therapeutic challenges due to their potential to replace lost neural and glial cells and support tissue repair and connectivity restoration ^17,59,60^.

Historically, astrocytes have been considered detrimental post-SCI due to their role in glial scar formation, and consequently, early strategies aimed to inhibit or deplete astrocytes to reduce scarring and promote regeneration ^61^. However, subsequent studies have revealed the supportive role of astrocytes in tissue repair, including their ability to promote axonal growth following injury ^62–64^. Particularly, transplantation of neonatally-derived cortical astrocytes into hemisected adult rat spinal cords has shown that these cells can survive, integrate into host tissue, and promote axonal regeneration at the lesion site ^65,66^. Similarly, transplantation of immature astrocytes into brain injuries has also shown improved recovery outcomes ^67,68^. These findings highlight that the specific phenotypes and localization of astrocytes are key determinants of a permissive environment for axonal regeneration ^61^, underscoring their multifaceted roles in neuroprotection and repair mechanisms ^23,69^.

### hAPCs as Candidates for Transplantation in Spinal Cord Injury

To harness the regenerative potential of immature astrocytes while leveraging the advantages of iPSCs, we generated human iPSC-derived astrocyte progenitor cells (hAPCs) to evaluate their integration into host spinal cord tissue, to optimize their future therapeutic use in SCI. Using two iPSC lines (WTC11 and KOLF2.1J), we tested a robust differentiation protocol to generate hAPC populations ^40^.

Consistent with previous studies ^40,49–52^, our *in vitro* differentiation protocol yielded astrocyte progenitor cells with high efficiency across both lines (Fig. 1). Immunocytochemical analysis of hAPCs at day 28 revealed strong expression of the glial-restricted progenitor marker A2B5 and the astrocyte progenitor marker S100β, with low staining for the neuronal precursor marker doublecortin (DCX), indicating a predominantly glial progenitor identity (Fig. 1C a-b, 1D a-b).

RT-qPCR confirmed elevated transcript levels of *ST8SIA1* and *S100β* in both cell lines, supporting their glial progenitor identity. In contrast, *GFAP*, *MAP2*, and *SOX2* expression was low in the iPSC-derived hAPCs, suggesting some cell heterogeneity, which may enhance cell survival and integration post-transplantation.

Although minor differences in differentiation capacity and marker expression were observed between the two iPSC lines, both demonstrated consistent glial lineage commitment. Importantly, hAPCs can be expanded, maintained, and genetically modified, and offer the potential for autologous transplantation. These characteristics position hAPCs as a promising candidate for future cell-based therapies aimed at spinal cord repair.

### *In vivo* assessment of short- and long-term survival, and phenotypic characterization

One of the key challenges we aimed to explore in transplanting astrocyte progenitors in the intact rodent spinal cord was their ability to survive. Previous studies have already suggested that transplantation of less mature human fetal-derived and hiPSC-derived glial precursor cells results in much improved survival, most likely because of their capacity to proliferate *in vivo* after transplant ^70–73^. Accordingly, here we showed that, on day 7 and 21 post-grafting, the WTC11- hAPCs-GFP^+^ and KOLF2.1J-hAPCs-GFP^+^ grafted cells survived in the grey matter of the spinal cord (Fig. 2 E a-c, F a-c). However, a decrease in the number of transplanted cells was found at both time points (7- and 21-day post-transplantation) (Fig. 2G). This may result from most cells either failing to engraft at the transplantation site, being lost during surgery, or not surviving beyond the first week after transplantation. A similar outcome was reported by Haidet-Phillips et al. ^73^, where <5% of the transplanted cells remained viable two weeks post-transplantation. Importantly, here, no significant differences in percentage of survival were noted within each line WTC11 and KOLF2.1J after transplantation at any of the time points examined (7- and 21-day post-transplantation).

Another goal of our study was to evaluate how the transplanted cells change between *in vitro* culture conditions and the *in vivo* environment of the intact spinal cord. On day 7 post-grafting, the WTC11-hAPCs-GFP^+^ tested positive for the A2B5 marker, confirming the preservation of the glial progenitor phenotype (Fig. 3A), even with a low expression of the GFAP marker. Conversely, on day 21 post-grafting, the transplanted cells resulted strongly positive for the astrocytic marker GFAP (Fig. 3B), indicating the change of phenotype. So, as our *in vitro* data anticipated (Fig. 1C-D), these findings suggested that the transplanted hAPCs were glial-committed and underwent phenotypic transformation, differentiating into mature astrocytes. Our data aligns perfectly with the previous studies showing that the hiPSC-derived astrocyte progenitors grafted in the C6 survived and differentiated into mature astrocytes ^73^.

Another important parameter to consider was the cell localization. Both the WTC11-hAPCs-GFP^+^ and KOLF2.1J-hAPCs-GFP^+^ transplanted cells exhibited a consistent distribution across all the regions and time points analyzed (7- and 21-day post-transplantation) (Fig.2). This reflected previous studies conducted in our lab, where the location of the transplanted cells within the intact spinal cord remained stable for up to 15 months ^16^.

### Lack of adverse outcomes

The lack of negative outcomes, such as tumors, uncontrolled proliferation, and localization in undesired areas, is also worth consideration.

Considering that the hAPCs used for the transplant were derived from hiPSCs, there was a significant risk of uncontrolled proliferation and tumor formation, largely attributed to undifferentiated cells ^74–76^.

Although both WTC11-derived hAPCs and KOLF2.1J-derived hAPCs exhibited low-level expression of the pluripotency gene *SOX2* (Fig. 1E), no tumor formation was observed at either 7- or 21-day post-transplantation. Indeed, as expected, the number of transplanted hAPCs decreased at 7- and 21-days post-grafting compared to day 0 (Fig. 2), indicating the absence of uncontrolled proliferation.

As for the localization, the hAPCs were observed relatively near the injection site, at either 7- or 21-day post-transplantation. This can be the result of the delivery method used here. While less invasive transplantation methods (i.e., intraventricular injection) are available ^77,78^, we chose the direct injection approach based on prior studies conducted by our lab ^16^. This finding highlights the safety of these cells for potential future transplant applications.

### Challenges and future directions

Although this study demonstrated the potential benefits and safety of transplanting hiPSC-derived astrocytic progenitor cells into the intact spinal cord, numerous questions and future challenges remain open. First, a necessary future study should address the transplantation of hAPCs after cervical spinal cord injury (C3-4) to assess the anatomical integration of donor cells and restoration of the phrenic pathways.

Furthermore, several recent studies have found a growing appreciation for the role of the glutamatergic spinal V2a neurons in contributing to respiratory plasticity following SCI ^79^. Notably, some groups revealed that transplanting stem cell-derived, engineered V2a neurons into the injured cervical spinal cord enhanced recovery beyond using less-defined mixtures of neuronal and glial progenitor cells alone ^80,81^. Building on this important knowledge, one future challenge could be the combined transplantation of the hiPSCs-derived APCs with the hiPSCs-derived V2a. Due to the uniform and glial-differentiation commitment of hAPCs, transplanting patient-derived hiPSC astrocyte progenitors into the spinal cord could be a viable approach for *in vivo* disease modeling and therapeutic applications.

## CONCLUSION

Here, we demonstrate the reliable and efficient generation of human astrocyte progenitor cells (hAPCs) from two iPSC lines (WTC11 and KOLF2.1J) using straightforward differentiation protocols. Our *in vivo* data provide an in-depth analysis of the short (7 days) and long (21 days) outcomes of transplanted hAPCs into intact rodent spinal cord, highlighting their ability to survive and become mature astrocytes, without inducing harmful effects. Particularly, we observed that the 2 iPSC lines used behaved similarly both *in vitro* and *in vivo* concerning differentiation, survival, and migration.

These long-term findings suggest that this defined population of hAPCs holds significant potential as a component of therapeutic strategies for SCI. Future advances will provide even greater studies on the phenotypic fate of hAPCs transplanted in the cavity after SCI.

## ACKNOWLEDGMENTS

AN was supported by the Craig H. Neilsen Foundation (430069) and the Christopher Reeve Foundation (284314). The William P. Snyder, III Chair Endowment supported IF. LQ is supported by National Institutes of Health (R01NS115977), a research grant received from ALS FINDING a CURE and HOP on A CURE, and Alzheimer’s Association research grant (24AARG-D-1191264). We thank Drs. Angelo Lepore and Brigid Jensen from Thomas Jefferson University (Philadelphia) for the helpful comments during the project’s development. We thank Julien Bouyer from Drexel University for the confocal microscope training.

## AUTHOR CONTRIBUTIONS

Dr. Niceforo designed and conducted all the experiments, analyzed the data, and wrote the manuscript. Dr. Jin helped with the animal surgeries, the interpretation of the *in vivo* data, and revised the manuscript. Skandha Ramakrishnan produced the lentivirus for the study and wrote the material and methods section related to the virus. Maurice Lindner-Jackson helped with animal care.

Drs. Fischer and Qiang, designed, supervised, and assisted with experiments, data interpretation, and analysis, and revised the manuscript.

All authors read and approved the final version of the submitted manuscript.

## DECLARATION OF INTERESTS

The authors declare no competing interests.

## Notes

### Competing Interest Statement

The authors have declared no competing interest.

